# Widespread detection of Polyethylene glycol reveals chronic human exposure through pharmaceutical drugs

**DOI:** 10.64898/2026.09.11.750999

**Authors:** Harsha Gouda, Patricia Kelly, Luke Whiley, Daniel Gómez-Pérez, Anna Mascellani Bergo, Wilhan Donizete Gonçalves Nunes, Elena Chekmeneva, Ada H.Y. Yuen, Mark David, Shona McKirdy, Andrew Nelson, Darren L. Smith, Haoqi Nina Zhao, Jeong In Seo, Edward J. Mathias, Gillian Farrell, Toon Scheurink, Michael Strobel, Helena Mannochio-Russo, Konstantinos Gkikas, Mingxun Wang, Jaroslav Havlik, Christopher Quince, Zoltan Takats, Matthew Lewis, Konstantinos Gerasimidis, Nicholas J. W. Rattray, Pieter C. Dorrestein

## Abstract

Polyethylene glycol (PEG) is a synthetic polymer ubiquitous in pharmaceuticals, personal care products, food additives, and industrial manufacturing. Despite its widespread use and potential importance as an exposure chemical, the prevalence of PEG exposure and its excretion in human populations remain largely uncharacterized. Moreover, PEG detected in human biofluids is frequently assumed to arise from analytical contamination during sample preparation, potentially obscuring its contribution to the human exposome and confounding metabolic phenotyping studies. Here, we show that PEG as a component of the human xenobiotic exposome and is further metabolized into PEG hydoxy acid and diacid metabolites in humans. We further find that PEG exposure is associated with alterations in microbiome composition and short-chain fatty acid metabolism, suggesting its biological impact of exposure. We identify PEG exposure in approximately 2.3% of publicly available metabolomics data files and provide a reusable 85,484 candidate PEG and PEGylated MS/MS spectral library for future use for the metabolomic community. Together, these findings establish PEG in human biofluids can reflect genuine exposure, and that PEG exposure is neither metabolically inert nor biologically silent.

## Introduction

Population health studies are increasingly probing the human metabolome to understand the molecular underpinnings of health and disease. Advances in metabolic phenotyping technologies and computational biology have enabled the generation of molecular profiles from large epidemiological cohorts that were previously the domain of genome sequencing and analysis.^1–3^ Beyond the metabolome, the technologies used inevitably capture instances of chemical exposure, such as those from pharmaceutical drugs, food and food additives, personal healthcare chemicals, and environmental pollutions.^4–6^ These exposures are now considered complementary to the genome with regard to their influence on health and disease status.^1,7^ In the meanwhile, detection of these chemicals can sometimes interfere with the measurement of the internal metabolome. To distinguish the exposure and the downstream metabolic changes caused by them, it is essential to annotate chemical exposure in the investigation of the metabolome.^8,9^

Polyethylene glycol (PEG) is a polyether composed of repeating sub-units of ethylene oxide widely present in a number of common over-the-counter and prescription drug formulations, food products, supplements, and personal care products.^4,10,11^ PEG compounds are presumed to be chemically inert, minimally absorbed, and excreted largely unchanged as detected in biological fluids.^12–14^ PEG-based laxatives are routinely administered to clinical patient cohorts and healthy individuals.^15,16^ PEG formulations are available not only by prescription but also in many over-the-counter drugs such as MiraLAX^®^, often marketed as safe and gentle remedies for occasional constipation.^17,18^ This accessibility, coupled with their widespread use, enables PEG chemical exposure in both clinical and general populations. Importantly, individuals may self-administer these agents without disclosure, leading to inadvertent inclusion of PEG-containing samples in epidemiological studies. Polyethylene glycol residues persist through standard sample preparation workflows, particularly in mass spectrometry.^19,20^ Low molecular weight PEG mixtures (e.g., PEG400) are also notorious within analytical chemistry as ubiquitous contaminants of sample collection containers, plastic labware, plastic tubing, LC column manufacture processes, detergents and common analytical reagents. Due to its polymeric nature, it is easily recognised in liquid chromatography mass spectrometry (LC-MS) assays as an envelope of repeating signals separated by 44.0262 Daltons. These patterns can have an obscuring or suppressive effect on the metabolic phenotype, representing a risk to the validity of data derived from PEG-containing biofluids.^21^

Occasional examples of PEG toxicity have been reported.^22^ The World Health Organisation has previously set an acceptable daily intake of 10 mg/kg (body weight) of PEG polymers.^23^ Emerging evidence suggests that its osmotic activity and interaction with the gut environment may exert subtle yet measurable biological effects.^24–26^ Studies in animal models and human cohorts have shown that PEG-based bowel preparations can transiently alter gut microbial composition and reduce bacterial diversity, possibly through mechanical clearance and osmotic stress.^13,24,27^ However, the direct effect of PEG on gut microbiome composition and function remains to be investigated. These findings indicate that PEG exposure may not only interfere with analytical measurements but also modulate the underlying biological milieu captured in biofluids.^28^ Understanding these dual effects is essential for accurately interpreting metabolomic and microbiome data derived from samples collected in clinical cohorts, particularly after detected PEG exposure.

In this study, we report that the detection of polyethylene glycol in metabolomics analysis of biological fluids arises may also from prevalent exposure through pharmaceutical medication rather than analytical contamination from sample preparation protocols. Through LC-MS/MS and ^1^H NMR spectroscopy, we identify concordance in PEG detection in samples with laxative intake in clinical cohorts of individuals with Crohn’s disease. While presumed to be inert, we show that PEG exposure has biological effects. PEG directly alters fecal microbiome beta diversity, short-chain fatty acid production, and microbial community composition. Subsequently, PEG can be oxidized to hydroxy acid (PEG-COOH) and diacid (PEG-2COOH) following PEG exposure through an over-the-counter gel capsule, as detected in blood and urine samples. Using public metabolomics data, we provide a reusable mass spectrometry library of candidate PEG and PEGylated ions for detection of PEG-related metabolites in future metabolomic studies.

## Results

### Laxative use is associated with a strong PEG signal in LC-MS/MS data

We detect polyethylene glycol in fecal metabolomic sample analyses of various cohorts in our laboratory. Some example cohorts include samples from Pitt Hopkins syndrome^29^, Rheumatoid arthritis^30^, Parkinson’s disease^31^, Crohn’s patients (iPENS cohort)^32^, and many more, which highlighted the widespread exposure across many clinical cohorts and disease states. As an example, we assessed the presence of polyethylene glycol (PEG) in stool samples from the iPENS cohort (n=144 samples; 11 PEG+ from 5 participants) (**Figure 1**).^32^ PEG+ samples showed approximately 1.5-fold higher total ion count than PEG-samples (median 2.10 × 10¹⁰ vs 1.27 × 10¹⁰; p < 0.001), and PEG-derived ions accounted for a median 31% of the total ion current in those samples versus 0.07% elsewhere (**Figure 1**). The laxative containing sample displayed ∼200-fold difference in PEG abundance between groups, a continuous signal envelope spanning a broad retention time, which was missing in samples without laxative exposure, indicating that this signal originates predominantly from the sample rather than the analytical sample preparation workflow.

**Figure 1:**
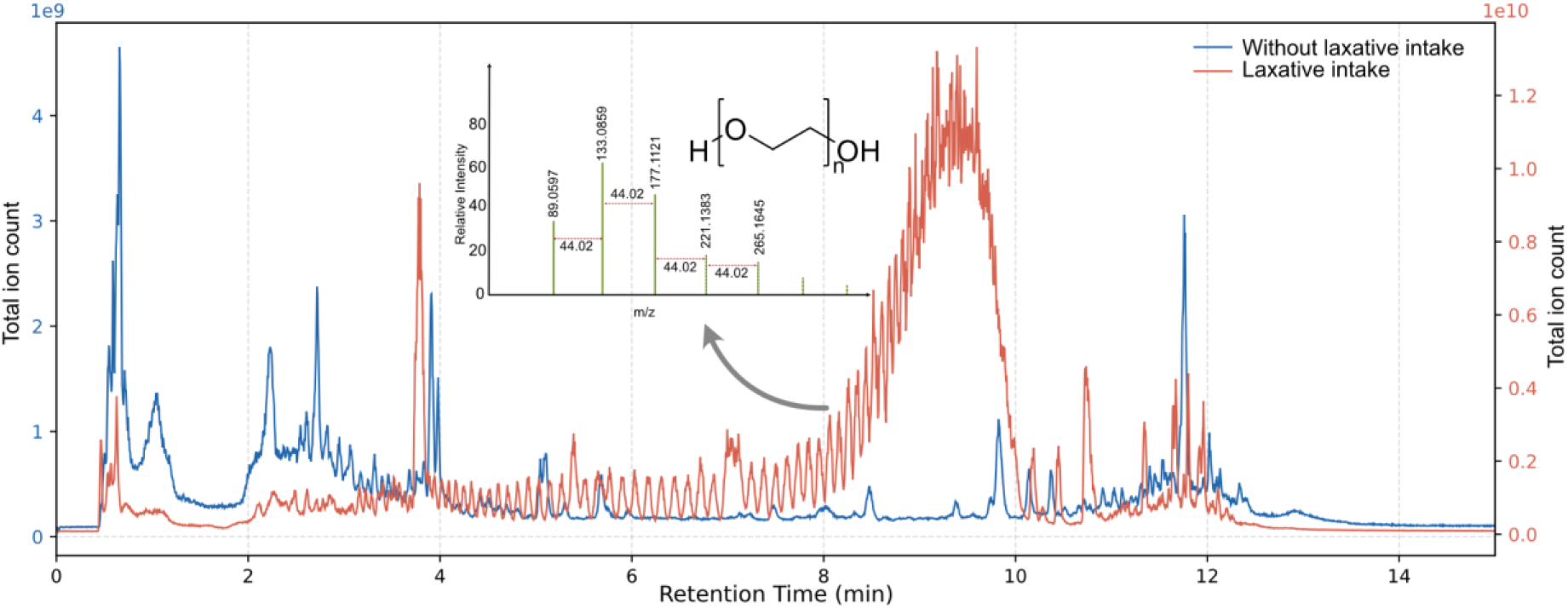
Chromatographic and spectral characterization of Laxative exposure in LC-MS samples: Total ion chromatograms (TIC) comparing stool samples with (orange) and without (blue) documented laxative intake. Inset plot shows MS/MS fragmentation pattern of PEG residues, with successive MS2 fragments separated by 44.02 Da.

MS/MS fragmentation of PEG precursor ions yields a diagnostic oxonium ion series at *m/z* 89.0597, 133.0859, 177.1121, 221.1383, and 265.1645, with successive ions separated by 44.0262 Da, corresponding to the exact mass of the ethylene oxide repeating unit (C₂ H₄ O; **Figure 1a, inset**). This regular 44 Da ladder is the hallmark fragmentation pattern of PEG polymers and confirms the PEG exposure from laxatives in iPENS stool samples. PEG ions were detected across a precursor mass range of *m/z* ∼150–1500 and eluted across the full chromatographic window from 2 to 10 minutes, confirming that PEG is not confined to a discrete chromatographic region but instead pervades the run over a long retention time window. The plot revealed a structured diagonal banding pattern in which higher *m/z* precursors eluted at progressively later retention times. PEG standards for varying average molecular weight PEG400, PEG1000, PEG3350 and PEG8000 showed a chain-length-dependent reversed-phase retention of PEG oligomers, spanning from low-mass short-chain oligomers eluting early to high-mass long-chain species eluting later in the chromatogram (**Figure S1**). Similarly, analysing PEG1000 on a trapped ion mobility-mass spectrometer, we observed a linear dependent increase in collision cross section (CCS) values for each additional ethylene oxide unit across multiple detected adduct forms (**Figure S2**).

### Detection of PEG signals by ^1^H NMR spectroscopy in iPENS cohort

As mentioned above, 1H NMR data was collected for iPENS cohort to investigate how the presence of PEG affects the NMR metabolomics data. Within ¹H NMR spectroscopy, PEG produces a characteristic singlet resonance at δ 3.65–3.70 ppm in aqueous fecal extracts, corresponding to the methylene protons (–OCH₂ CH₂ –) of the ethylene oxide repeating unit^33^. This signal appears as a broad, prominent peak, readily identifiable against the background of fecal metabolites (**Figure 2a**). Unlike in LC-MS/MS, where PEG contamination substantially distorts chromatographic profiles and suppresses metabolite detection, NMR-based fecal metabolomics is considerably more tolerant of PEG presence. The PEG signal does not interfere with spectral acquisition; however, it partially overlaps with resonances of other metabolites in the approximately 3.3–3.8 ppm region. Human fecal ¹H NMR spectra commonly contain signals from short-chain fatty acids, amino acids, sugars, organic acids, and amines. Depending on the exact position and width of the PEG peak, the most affected metabolites are expected to include compounds with resonances in this region, such as glucose and other carbohydrates, glycerol, glycine, betaine, choline-containing compounds, taurine, myo-inositol, and the α-proton resonances of several amino acids, including alanine, serine, aspartate, and related signals (**Figure 2b**). Importantly, because most of these metabolites also resonate in other spectral regions, the relevant biochemical information can still be recovered from unaffected spectral regions, thereby reducing the analytical impact of PEG contamination in NMR-based analyses. NMR spectroscopy additionally offers molecular weight estimation of detected PEG. Analysis of ¹H chemical shift characteristics and signal linewidth can be used to distinguish PEG grades^33^. Based on these approaches, the PEG detected in iPENS samples was consistent with PEG 3350.

**Figure 2.**
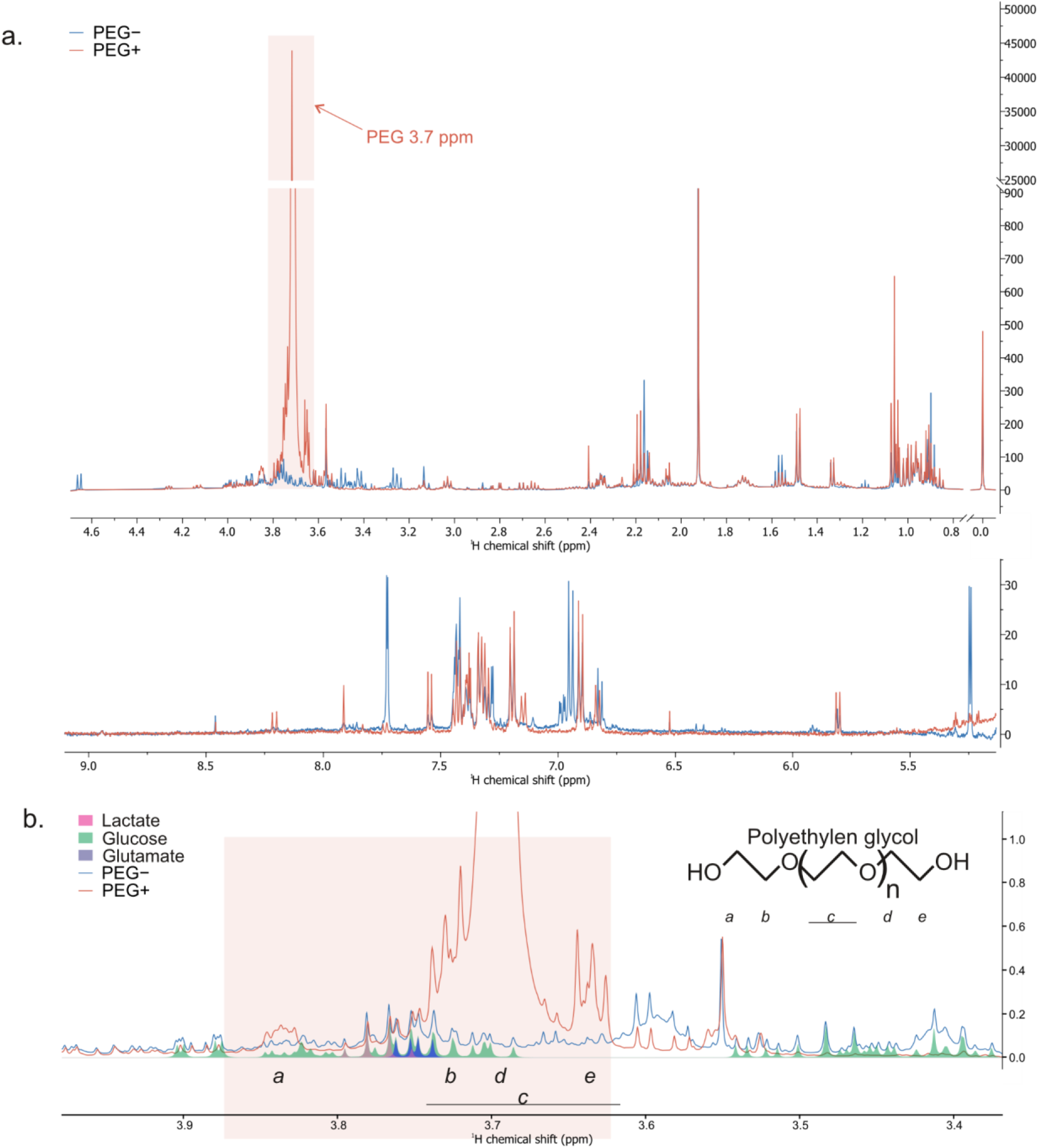
¹H NMR spectra of fecal samples with and without polyethylene glycol (PEG) signal. (a) Full metabolite region (0.5–5.0 ppm, top) and aromatic region (5.0–9.0 ppm, bottom). The orange-shaded area highlights the PEG region (3.59–3.84 ppm). (b) Expanded view of the PEG region, where the dominant PEG signal in sample IPENS012A (PEG+, red) is evident against the sample IPENS901D (PEG-, blue); Spectra were acquired at 500 MHz using a 1D NOESY sequence with presaturation water suppression.

To directly compare the two platforms, NMR and LC-MS/MS detection were performed independently and subsequently cross-validated. LC-MS/MS provided a binary classification (PEG+/PEG-) based on detection of the diagnostic oxonium ion series (*m/z* 89, 133, 177, 221, 265), while NMR provided a semi-quantitative measure of PEG load via the integrated intensity of the δ 3.65–3.70 ppm bin. As expected, two complementary methods showed concordance across all iPENS samples (100% agreement; **Figure S3**). PEG+ samples identified by LC-MS/MS displayed NMR signal intensities above the classification threshold, while PEG-samples clustered tightly below this threshold (**Figure S3**). Longitudinal profiling revealed that PEG exposure was persistent across multiple time points in a subset of subjects, whereas others showed transient or absent PEG signal, consistent with intermittent laxative use (**Figure S3**).

### PEG exposure alters microbiome composition and short-chain fatty acid metabolism

To determine whether PEG exposure through laxative intake was associated with alterations in gut microbial metabolism, fecal short-chain fatty acids (SCFAs) were quantified in 1,014 iPENS stool samples classified as PEG+ or PEG-as detected by the LC-MS/MS and NMR spectroscopy. Multivariate analysis of dry-weight normalised SCFA profiles demonstrated significant differences according to PEG exposure (PERMANOVA, F = 14.1, R2=0.015, p<0.0001). After accounting for repeated sampling from the same individual, PEG exposure remained significantly associated with overall SCFA composition (F = 4.2, R2=0.0047, p=0.0025), indicating that PEG exposure explained a small but reproducible proportion of the overall variation in fecal SCFA composition. Comparison of individual SCFAs demonstrated reduced concentrations of several SCFAs in PEG+ samples. Significant decreases were observed for all three major short-chain fatty acids (acetate, propionate, and butyrate) in PEG+ samples (**Figure 3a**). Significant changes in isobutyrate, 2-methylbutyrate, isocaproate, caproate, and caprylate were also observed.

**Figure 3.**
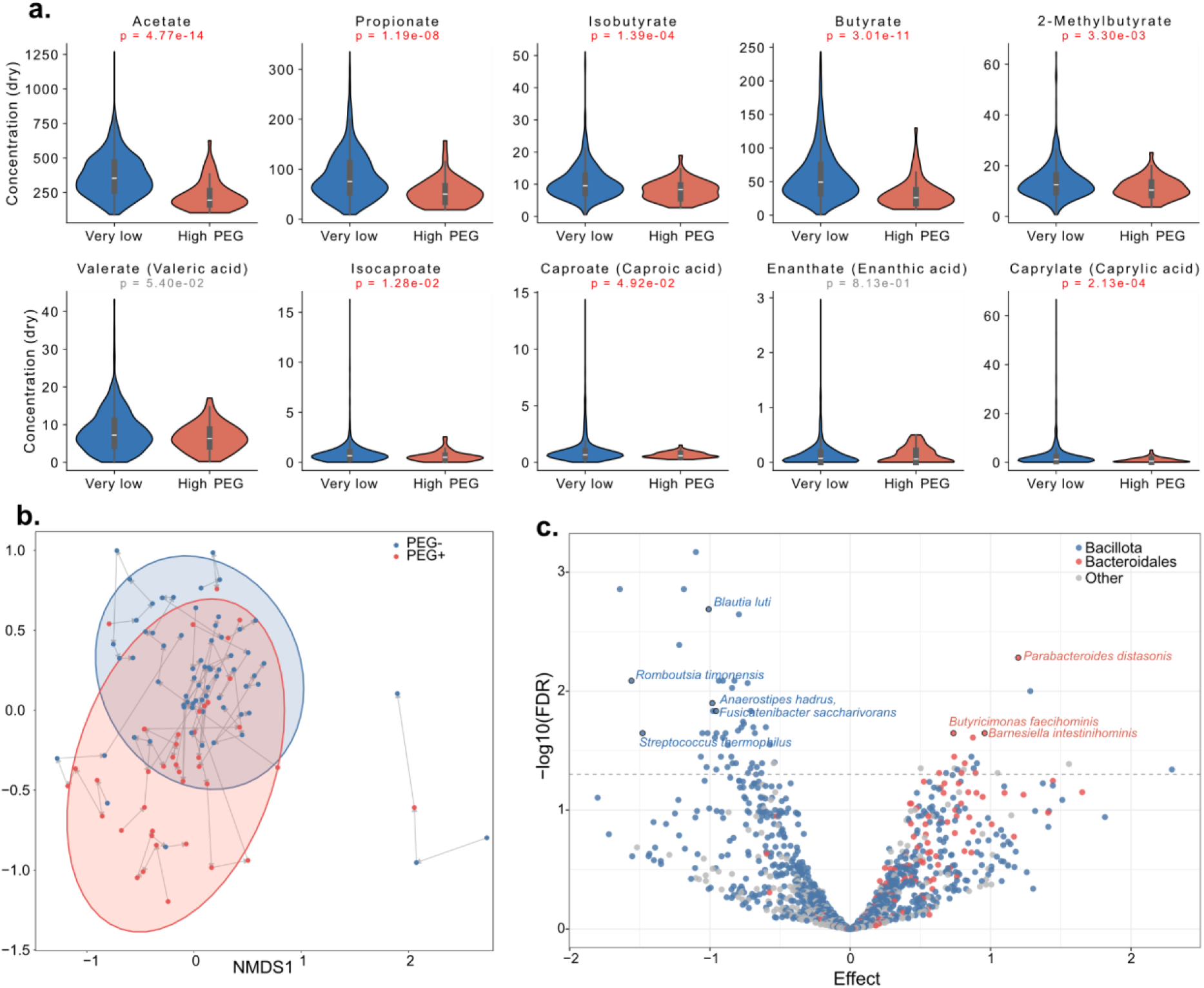
Polyethylene glycol (PEG) exposure is associated with altered fecal short-chain fatty acid (SCFA) concentrations and gut microbiome composition. (a) Fecal short-chain fatty acid profiles normalized to dry fecal mass in PEG+ and PEG-samples. (b) Non-metric multidimensional scaling (NMDS) ordination of gut microbial community composition from species-level MAG relative abundances comparing PEG+ (red) and PEG-(blue) stool samples. Arrows connect multiple samples from the same patient ordered by time point (n=109). (c) Differential abundance analysis of species-level MAGs associated with PEG exposure. Positive effect sizes indicate taxa enriched in PEG+ samples, whereas negative effect sizes indicate taxa depleted in PEG+ samples. MAGs are colored by taxonomy: Bacillota (blue), Bacteroidales (red) and other (grey).

To investigate whether PEG exposure was associated with changes in gut microbial community structure, Bray-Curtis beta diversity was assessed from representative species-level metagenome-assembled genome (MAG) abundance profiles for iPENS patients with both PEG+ and PEG-samples (n=16). Non-metric multidimensional scaling (NMDS) of MAG relative abundances demonstrated a shift in overall microbial community composition associated with PEG exposure (PERMANOVA, R2=0.0597, p<0.001; **Figure 3b**). The same pattern was recovered by read-based taxonomic profiling with MetaPhlAn 4 (*R*² = 0.0462, *p* < 0.001). Differential abundance analysis at the MAG level using 824 samples from 112 patients identified 69 significantly differentially abundant taxa (21 enriched in PEG+, 48 enriched in PEG-; FDR < 0.05), revealing a taxonomically structured and asymmetric response (**Figure 3c**). Depletion in PEG+ samples was almost exclusively confined to the phylum Bacillota, predominantly class Clostridia and family Lachnospiraceae, including the SCFA-producing genera *Blautia_A*, *Anaerostipes*, *Agathobacter*, and *Fusicatenibacter*. In contrast, taxa enriched in PEG+ samples were more phylogenetically diverse and dominated by the phylum Bacteroidota, including *Odoribacter*, *Butyricimonas*, *Alistipes*, *Parabacteroides distasonis*, and *Barnesiella*, together with members of Pseudomonadota (*Sutterella*, *Parasutterella*) and *Desulfovibrio* (Desulfobacterota). The concurrent loss of butyrogenic Lachnospiraceae and expansion of Bacteroidia and Proteobacteria is consistent with the reduced fecal SCFA concentrations observed above (**Figure 3a**), suggesting that PEG exposure shifts the community away from SCFA-producing Clostridia toward osmotically tolerant, bile-resistant taxa.

### Human PEG metabolism following medication exposure

Detection of PEG in blood within 4 h of oral medication has previously been reported.^34^ In the AIRWAVE study cohort, we observed the in vivo oxidation of PEG to diacid (PEG-2COOH) and hydroxy acid (PEG-COOH) metabolites, alongside the PEG parent ion in the urinary metabolomic profiles (**Figure 4a-b**).^35^ These forms of oxidized PEG metabolites have previously been reported as metabolites of PEG in mammalian studies.^36–38^ We observed a positive intra-sample correlation (Pearson r =0.92) between the PEG (n8, [M+NH₄]⁺) ion and its oxidized hydroxyacid metabolite PEG(n8)-COOH, suggesting a relationship between exposure to PEG(n8) and its metabolic product (**Figure 4c**).

**Figure 4:**
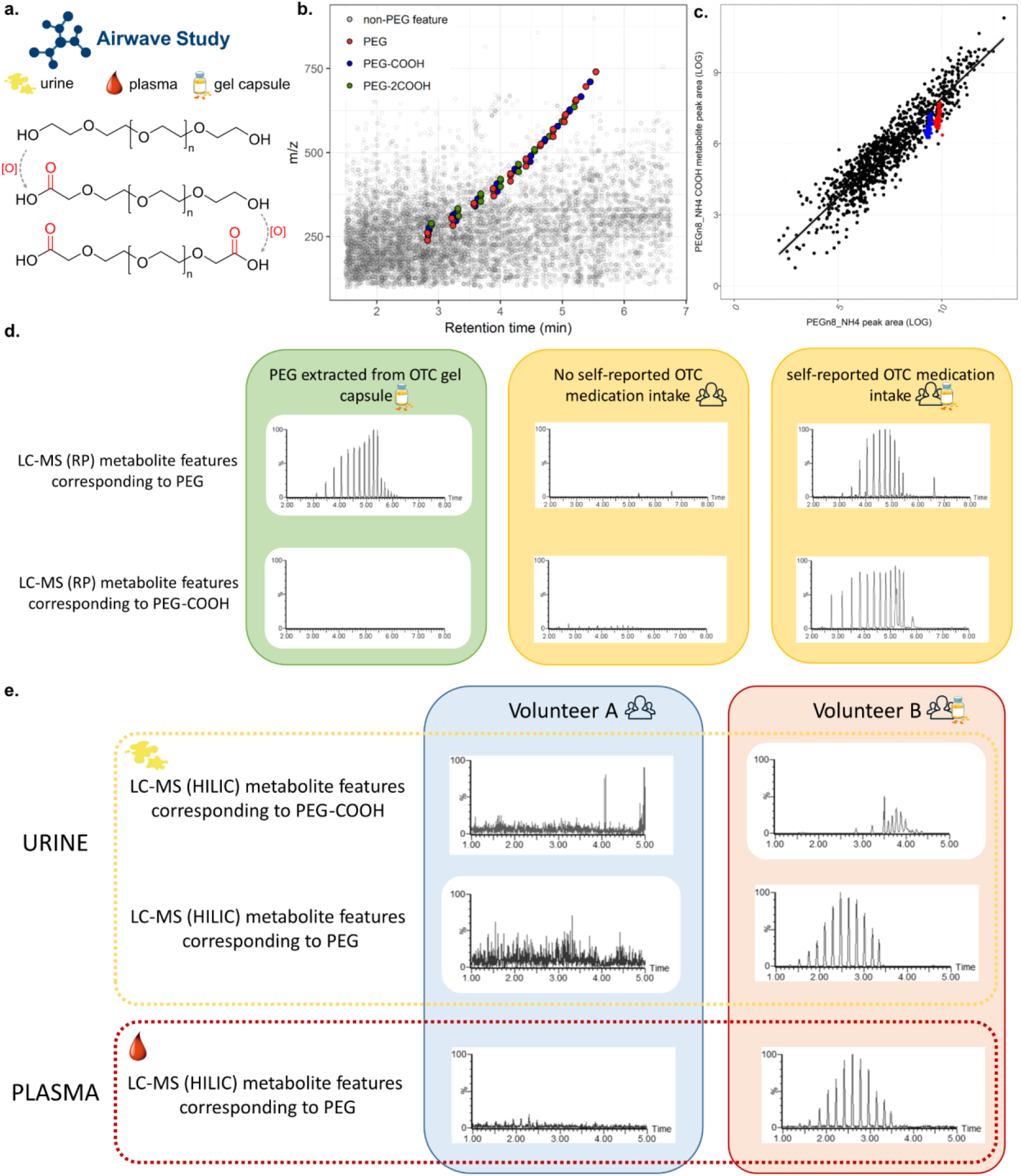
Detection of PEG oxidation metabolites following medication intake in AIRWAVE cohort. (a) Urine and plasma samples from AIRWAVE Health Management cohort were used to detect metabolites of PEG oxidation (b) An example urine metabolomic profile of reversed phase analysis with PEG metabolites highlighted, PEG (red), PEG-COOH (blue), PEG-2COOH (green) and non-PEG features (grey). (c) Extracted log peak area correlation between PEG (n8, [M+NH₄]⁺) and PEG-COOH (n8, [M+NH₄]⁺) in AIRWAVE cohort (n=990). Blue and red points (n=96 each) represent external and internal quality control samples acquired during analysis. (d) Extracted ion chromatogram of PEG and PEG-COOH (n5-16) over the counter gel capsule extraction, urinary samples with non-self reported and self reported gel capsule intake. (e) Extracted ion chromatogram profiles of PEG and PEG-COOH (n5-16) from plasma and urine biofluids in volunteers in the AIRWAVE cohort.

To characterize the biotransformation of PEG to PEG hydroxy acid metabolite following exposure, we compared mass spectrometry profiles of the over-the-counter gel capsule (Nurofen Express)^39^ medication containing PEG, and urine samples from individuals who self-reported the consumption of the over—the-counter gel containing pharmaceuticals (**Figure 4d**). We observed that the gel capsule contained only PEG, with no PEG-COOH metabolite. The mass spectrometry profile for the control urine samples contained low, background traces of PEG and PEG-COOH. In the urine sample collected following the self-reported consumption of an over-the-counter gel capsule, metabolite profiles contained both PEG and PEG-COOH at high concentrations, suggesting absorption, metabolism, and detection in urine samples (**Figure 4d**). In addition to the urine samples, we analyzed plasma samples collected during the same study visit. Participants with high observable levels of PEG and PEG-COOH in their urine also had high observable levels of PEG detected in plasma. Participants with urine that contained either low levels of PEG and PEG-COOH or contained no detectable trace of PEG/PEG-COOH also had low levels or no detectable trace of PEG in plasma (**Figure 4e**). The metabolite PEG-COOH was not detected in any of the plasma samples in this cohort. PEG and PEG-COOH signals in urine were adjusted for urine volume and dilution effect by performing a ratio with quantification values for creatinine (acquired by NMR). Data were subsequently LOG-transformed. Pearson correlation analysis resulted in a positive correlation (r = 0.32, *p*=<2.2e^-16^), suggesting a significant relationship between circulating plasma PEG and excreted urinary PEG-COOH metabolite (**Figure S4**).

Metabolomics analysis of the over-the-counter drugs available in the United States, such as Tylenol and Ibuprofen that are two of the most frequently consumed drugs (**Figure S5**),^40^ showed the ubiquitous detection of PEG ions in two of the most consumed over-the-counter medications, consistent with PEG being a widely used pharmaceutical excipient and tablet coating agent. We also identified PEG polymer signatures in a structurally distinct class of topical and transdermal drug products, such as menthol pain-relieving gel (brand: Mineral ice), clotrimazole cream (brand: Lotrimin), and Diclofenac dimethylammonium (brand: Voltaren Emugel 2%). We were also able to detect the PEG molecular ions across various food and personal care products (**Figure S6**) that contribute to the human exposome. For example, long-chain PEG ions were detected with the highest prevalence in sun lotion, facial cleanser, lotion and moisturizer, sunscreen, and serums.^6,41,42^

### Mass-spectrometry (MS2) library for PEG annotation

To enable detection of PEG and PEG metabolites, we utilised the mass spectrometry query language (MassQL) to query for diagnostic oxonium ion series at *m/z* 89.0597, 133.0859, 177.1121, 221.1383, and 265.1645 across public metabolomics data,^43^ and identified widespread detection of polyethylene glycol across all MS/MS samples available in the public metabolomics repositories (GNPS/MassIVE, Metabolomics Workbench, and MetaboLights). We observed 4,085,996 PEG molecular ions detected across three repositories. These corresponded to 66,644 files in MassIVE (total files: 2,477,862), 4,305 files in Metabolomics Workbench (total files: 204,085) and 3,388 files in MetaboLights (total files: 538,621) (**Figure 5a**), constituting presence in 2.3% of total files in public metabolomics data repository and highlighting widespread detection across clinical cohorts of metabolomics data analysis. For each candidate precursor mass detected, theoretical chain length was calculated by solving for integer N = [(precursor mass × z - 106.063 - z × adduct) / 44.026] + 2. Among 85,854 unique precursor *m/z* values, we were able to assign the 13,352 ions to chain lengths associated with these PEG ions, with <10 ppm error, corresponding to neutral molecular weights of ∼263.163 to ∼1,488.9 Da. For adduct mass, ammonium, sodium, potassium, and protium masses were used to calculate the neutral mass of the PEG polymeric ions. The remaining ions were derived from metabolism of PEG, PEGylated molecules (examples include polyethoxylated tallow amines, covalently PEG-linked protein/aptamer/small molecule drugs, PEG-lipid conjugates as part of vaccine formulations) or ion forms (ISF’s, adducts, multimers) that were not considered.

**Figure 5:**
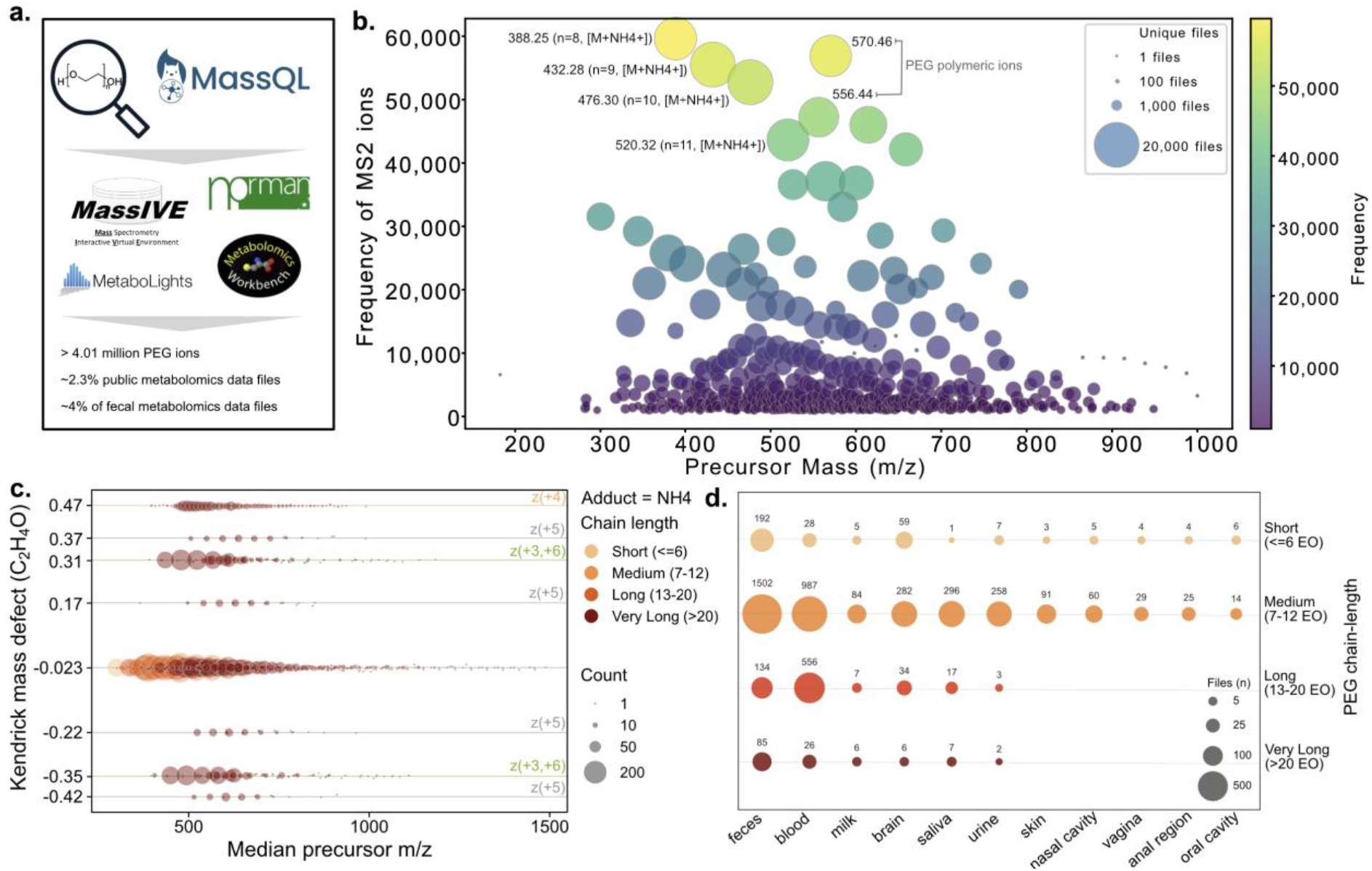
PEG detections across public metabolomics data repositories. (a) PEG structures were queried using MassQL workflow across three public metabolomics repositories (b) Bubble plot of precursor *m/z* versus MS2 ion detection frequency across the repository (size= unique files, color = frequency). (c) Kendrick mass defect (KMD) plot of candidate PEG libraries annotated with chain length, charge state and adduct form. (d) Annotated PEG spectral detection across anatomical UBERON body part sites.

The most frequently detected PEG ions across metabolomics repositories were the medium-chain [M+NH₄]⁺ series: PEG-8 (*m/z* 388.25, 59,617 detections), PEG-9 (*m/z* 432.28, 55,531 detections), PEG-10 (*m/z* 476.31, 52,750 detections), and PEG-11 (*m/z* 520.33, 43,641 detections), consistent with low-molecular-weight PEG400 (n=9) commonly used as pharmaceutical excipients (**Figure 5b**)^26^. Higher molecular weight PEG species (n=21-41) were detected as multiply charged [M+2NH₄]²⁺ and [M+3NH₄]³⁺ ions in the *m/z* range 335-688. Stratifying detected PEG ions by adducts revealed that NH4+ adducts accounted for the highest total detection frequency across the public repositories, with various charge states (**Figure 5c**). We further resolved their distribution in data from human biological sample types, using the available curated metadata files.^44^ The results highlighted that the PEG detection is not confined to a single biofluid or sample type but is instead broadly distributed across diverse biological matrices (**Figure 5d**). In addition to feces, blood and urine as previously described in the literature, PEG signal was further detected in saliva, skin, brain, human milk and nasal cavity. Across all body sites, the chain length distribution of detected PEG was dominated by medium-chain oligomers, though very long chain species, consistent with laxative-type PEG 3350, were disproportionately represented in fecal and blood samples, reflecting likely oral ingestion and systemic absorption routes. The candidate library PEG and PEG metabolites with 85,484 MS/MS spectra is now made available for public use using Global Natural Product Social Molecular Networking (GNPS) infrastructure.^45^

## Discussion

Analytical laboratories performing metabolic profiling have historically viewed PEG as a contaminant introduced by laboratory equipment, sample handling, or chromatographic solvents, and have routinely filtered these features from downstream analysis.^46^ Here we show that, in biological samples of the cohorts used in this study, PEG signal predominantly reflects genuine human exposure rather than analytical artifact. Across LC-MS/MS and ¹H NMR platforms, PEG detection was concordant with laxative intake in clinical cohorts, persisted longitudinally in line with intermittent use, and critically gave rise to oxidative metabolites that cannot originate from labware. These observations reposition PEG from a nuisance to be discarded into an informative and underappreciated component of the human chemical exposome.

The scale of exposure is substantial. PEG-associated ions were detected in approximately 2.3% of publicly available metabolomics files spanning GNPS/MassIVE, Metabolomics Workbench, and MetaboLights, and were distributed across diverse biological matrices: feces, blood, and urine as previously reported, but also saliva, skin, brain, human milk, and the nasal cavity, indicating that exposure is neither rare nor confined to the gut.^44,47^ This ubiquity is consistent with the pervasive use of PEG as a prescription drug formulations^10^, food products^4,11^, supplements, infant diapers, and cosmetic ingredients^6,36^, which we confirmed by detecting PEG across common over-the-counter medications, topical products, foods, and personal care items. Because the World Health Organization acceptable daily intake for PEG (10 mg/kg body weight) was established primarily on toxicological rather than exposome grounds, the frequency and multiplicity of exposure routes documented here argue that population-scale monitoring of PEG is warranted.^23^ To enable this, we provide a reusable MassQL-based MS/MS library of candidate PEG and PEGylated ions, allowing PEG exposure to be systematically annotated rather than silently removed in future phenotyping and epidemiological studies.^8,9^

PEG is not merely present but metabolically processed. In the AIRWAVE cohort, we observed in vivo oxidation of PEG to its hydroxy-acid (PEG-COOH) and diacid (PEG-2COOH) forms, consistent with prior reports of PEG oxidation by alcohol dehydrogenase in mammalian systems.^37,38^ The strong intra-sample correlation between PEG(n8) and its oxidized product (r = 0.92) links parent exposure to metabolic conversion, and the self-administration experiment provides direct causal support: the over-the-counter gel capsule contained only unmodified PEG, yet urine collected after ingestion contained both PEG and PEG-COOH at high abundance, while the capsule alone and control urine did not. Concordant PEG signal in matched plasma and urine (r = 0.32, p < 2.2 × 10⁻ ¹⁶) further demonstrates systemic absorption and renal excretion. Together, these data establish that orally ingested PEG is absorbed, biotransformed, and excreted a metabolic fate at odds with the long-standing assumption that PEG is chemically inert and passes through the body unchanged.^12,13^

Beyond its analytical footprint, PEG exposure was associated with a reproducible restructuring of the gut microbial community. Fecal short-chain fatty acid concentrations were significantly lower in PEG-exposed samples (PERMANOVA p < 0.0001; p = 0.0025 after accounting for repeated sampling), with reductions in acetate, propionate, and butyrate. At the species (MAG) level, this loss was mechanistically mirrored in the taxa affected: depletion was almost entirely confined to Bacillota of the class Clostridia, and specifically to butyrogenic Lachnospiraceae, including *Blautia*, *Anaerostipes hadrus*, *Fusicatenibacter saccharivorans*, and *Agathobacter* among the principal producers of the very SCFAs that declined. Conversely, PEG-exposed samples were enriched in bile-tolerant Bacteroidia (*Odoribacter*, *Butyricimonas*, *Alistipes*, *Parabacteroides distasonis*, *Barnesiella*), Pseudomonadota (*Sutterella*, *Parasutterella*), and the sulfate-reducing *Desulfovibrio*, a compositional signature consistent with an osmotic, dysbiotic perturbation. These findings align with prior evidence that PEG-based laxatives and osmotic stress transiently reduce bacterial diversity and can drive long-term community change, that PEG400 alters microbiota composition and host metabolism in mice, and that PEG modulates microbial bile-salt and lipid metabolism in the intestine.^24^ Chronic or repeated exposure that erodes SCFA-producing commensals is therefore a plausible, if still speculative, route by which PEG could exert biological effects beyond its intended osmotic action.

These biological findings carry direct methodological consequences. In LC-MS/MS, PEG signal is not confined to a discrete chromatographic region but pervades the entire analytical window, suppressing endogenous metabolite signals, driving retention-time drift, and reducing the number of detectable features; removal of PEG features alone can introduce confounding through uncorrected ion suppression. By contrast, ¹H NMR is comparatively tolerant: the broad PEG singlet at δ 3.6–3.9 ppm overlaps carbohydrate, amino acid, and choline resonances but is spectrally confined, allowing biochemical information to be recovered from unaffected regions and even permitting semi-quantitative estimation of PEG load and grade. This platform contrast, combined with the prevalence of exposure documented here, argues for treating PEG as a measured covariate rather than a discarded artifact. It also supports multi-platform study designs: a rapid ¹H NMR screen could flag PEG-containing samples before MS injection, avoiding both the ion suppression and cross-sample contamination in mass spectrometry.

It is not within the scope of this article to speculate on the long-term health effects of chronic exposure to PEG, however, it is apparent that this exposure to PEG should no longer be disregarded as a laboratory contamination and should be considered as part of the human exposome. Environmental exposome chemicals will influence our health status and are becoming part of the human metabolite phenotype. The work presented here also presents a strong case for changes in experimental design, demonstrating increased importance in the metabolite profiling of multiple biofluids collected simultaneously. As metabolite phenotyping is now frequently applied to large clinical and epidemiological sample cohorts, the acquisition of both samples and the raw data is both time-consuming and costly. Polymer contaminants are known to interfere with and suppress the signal of co-eluting analytes in LC-MS experiments; therefore, the high presence of PEG in biological samples has the potential to impact overall dataset quality. A reduction of PEG in samples designated for metabolite phenotyping would ensure the acquisition of analytical data that is of better quality and would be more appropriate for population phenotyping and retrospective data mining. As sample collection protocols are developed that are specific for metabolic phenotyping, the implementation of the restriction or substitution of products that are known to contain PEG for biofluid study participants in the lead up to sample collection may be beneficial to study design. An example that could be implemented is the inclusion of laxative intake as an exclusion criterion, or substitution of over–the-counter pharmaceuticals in gel capsules could be replaced with powder form alternatives that do not contain PEG.

## Author Contributions

H.G. and P.K. contributed equally to this work. P.K., G.F. and N.J.W.R designed the analytical strategy, performed data acquisition for iPENS fecal metabolomics data. H.G. performed LC-MS/MS analysis of iPENS samples, PEG standards, over-the-counter medications, food, and personal care products, and developed the MassQL query strategy and repository-scale search. K.Ger., K.Gk. and S.M. designed and conducted the iPENS clinical trial and coordinated sample collection, and contributed to iPENS sample processing. A.M.B. and J.H. performed ¹H NMR spectroscopy acquisition, spectral processing, and PEG signal quantification, and contributed to cross-platform comparison. L.W., E.C., A.H.Y.Y., M.D., M.L., and Z.T. designed and performed metabolic phenotyping of AIRWAVE urine and plasma samples, contributed the PEG oxidation metabolite analysis, and interpreted the biotransformation data. D.G.P. and C.Q. performed metagenomic assembly, MAG reconstruction, taxonomic annotation, and differential abundance analysis. W.D.G.N., H.M.-R., H.G., M.St., and M.W. developed the spectral library generation workflow, performed clustering and molecular networking-based filtering, and built the GNPS2 library infrastructure. H.N.Z., J.I.S., E.J.M., and T.S. contributed to data acquisition and analysis of PEG standards and drug formulations. H.G. wrote the original draft, and all authors reviewed, edited, and approved the final manuscript.

## Notes

The authors declare the following competing financial interest(s): P.C.D. is an advisor and holds equity in Cybele, Sirenas, and BileOmix, and he is a scientific co-founder, advisor, and holds equity to Ometa, Enveda, and Arome with prior approval by UC San Diego. P.C.D. consulted for DSM Animal Health in 2023. KGer has received funding for research and speakers fees from Nestle Health Sciences, Nutricia-Danone, AbbVie, Eli Lilly

## Acknowledgements

This project is supported by Crohn’s and Colitis Foundation (CCF, 1243263, 1243262 & 670398), Helmsley Charitable Trust and National Institute of Diabetes and Digestive and Kidney Disease (R01-DK136117) granted to P.C.D. H.G is supported by the American Heart Association (Grant: 26POST1559436) and Schmidt AI for Science Postdoctoral Fellow Programs. The AIRWAVE study was funded by the Home Office (Grant number 780-TETRA) with additional support from the National Institute for Health Research (NIHR), Imperial College Healthcare NHS Trust (ICHNT) and Imperial College Biomedical Research Centre (BRC) (Grant number BRC-P38084). The Airwave cohort is currently funded by the UKRI (grant award UKRI2539) and the NIHR HPRU in Radiation Threats & Hazards (NIHR-207424). Metabolic profiling of the urine and plasma samples from the AIRWAVE study carried out at the National Phenome Centre was supported by the Medical Research Council and National Institute for Health Research [grant number MC_PC_12025] and infrastructure support was provided by the National Institute for Health Research (NIHR) Imperial Biomedical Research Centre (BRC). NMR analysis was done using the METROFOOD-CZ Research Infrastructure (https://metrofood.cz), supported by the Ministry of Education, Youth and Sports of the Czech Republic (Project No. LM2023064). C.Q. and D.G.-P. acknowledge the support of the BBSRC funded EI ISP BBX011089/1 and BBS/E/ER/230002C. The iPENS study was funded by Nestle Health Sciences, J P Charitable Foundation, The Catherine McEwan Foundation, and the Crohn’s and Colitis Foundation. M.S. was supported by the National Institute of Allergy and Infectious Diseases under award no. 1F31AI200270-01. J.I.S. was supported by the National Research Foundation of Korea (NRF) (RS-2025-02373133).

## Data Availability

The candidate PEG and PEGylated ion MS/MS spectral library generated in this study is publicly available through the GNPS2 library infrastructure at https://library.gnps2.org/ (library identifier: GNPS2-PEG-MASSQL-PROPAGATED), and can be downloaded in .mgf or .json format.Untargeted LC-MS/MS data from the PEG standards [MSV000101530], and over-the-counter [MSV000099832], food [MSV000097542], and personal care product [MSV000095003] analyses have been deposited in the MassIVE repository. iPENS Crohn’s dietary intervention study and AIRWAVE Health Monitoring Study data are not publicly available owing to participant confidentiality and the terms of participant consent. Access requests can be made to Konstantinos Gerasimidis for iPENS data and Zoltan Takats for AIRWAVE study.

## Methods

### IPENS Clinical Trial

Ethical approval was obtained by the West of Scotland Research Ethics Committee 5 (19/WS/0163). A multicentre, prospective study was performed to identify dietary triggers of Crohn’s disease. Children and young adults with active Crohn’s disease (6-17 years) who were clinically responding to a 6–8-week course of exclusive enteral nutrition (EEN) were recruited from 11 paediatric hospitals across the UK. Patients were subsequently followed for 21 days during reintroduction of solid food, immediately after EEN completion, to explore associations between diet intake and recurrence of gut inflammation. Fecal samples were collected at set timepoints; days 3, 6, 9, 12, 15 and 21 following EEN completion. Samples were collected at patients’ homes, were immediately frozen in domestic freezers (−20°C) and transported frozen to the laboratory. After thawing, they were vortexed, aliquoted in Eppendorf tubes and stored in −80°C for downstream analysis. Following a second thaw cycle, they were aliquoted into a 96-well plate and prepared using an optimised dilute-and-shoot protocol. Metabolite extraction was performed using H₂ O as the extraction solvent with a 1:5 fecal: H₂ O dilution. The samples were mixed with a pipette to ensure solvent distribution, and the plate was submitted for UHPLC-MS analysis. ^13^C labelled tryptophan was used as an internal standard for assessing instrument stability and experimental conditions.

### CD-TREAT Clinical Trial

The study was approved by the West of Scotland Research Ethics Committee (reference 17/WS/0119). All participants and/or caregivers provided written consent. CD-TREAT was used as an induction therapy in adult and paediatric patients with active Crohn’s disease. After baseline assessments, patients were exclusively treated with CD-TREAT for up to 8 weeks, with all food provided. A total of 54 patients, including 27 adults (ages 18-57) and 27 paediatric (ages 8-17) CD patients were included. All patients with GIP detected in stool at timepoints B or C were excluded. Participants recorded their dietary intake daily. Stool, urine, and plasma samples were collected at baseline and at 4- and 8-week follow-up timepoints of the intervention. Treatment was discontinued if disease activity worsened at any time or if there was no improvement by week 4.

### LC-MS Analysis for iPENS cohort

Untargeted metabolomic analysis was performed on an ultra-high performance liquid chromatography (UHPLC) system (ThermoFisher Scientific) coupled to an Orbitrap Exploris 240 (ThermoFisher Scientific) mass spectrometer. Chromatographic separation was performed on a Vanquish Accucore C18 + UHPLC analytical column (ThermoScientific, 100 mm × 2.1 mm, 2.6 μM). Mobile phase A was composed of 99.9% water + 0.1% formic acid and mobile phase B was composed of 99.9% ACN + 0.1% formic acid. Electrospray ionisation (ESI) was used in positive mode (3900 V). The elution gradient used can be found in Supplementary Information Table S1. The source-dependent parameters were operated under the following conditions: sheath gas, 40 Arb; auxiliary gas, 10 Arb; sweep gas, 1 Arb; ion transfer tube temperature, 300 °C; vaporiser temperature, 280 °C. Instrument calibration was performed using Pierce^TM^ FlexMix^TM^ calibration solution (Thermo Scientific) and ran under vendor recommended settings. MS data collection was performed in data dependent acquisition mode (DDA) to give putative metabolite identification at MSI level 2^1^.

### Mass Spectrometry Data Processing

The acquired raw LC-MS/MS data were converted to .mzML format using MSConvert (ProteoWizard, Palo Alto, CA, USA), and the data was processed using MZmine4 version 4.3.0^2^. The mzML files were imported using the advanced import module with both MS1 and MS2 detection set to Factor of lowest signal, with a noise level set to 4 for MS1, and to 2.5 for MS2. The chromatogram construction was performed, where the minimum consecutive scans was set to 4, the minimum intensity for consecutive scans was set to 1.0E4, minimum absolute height set to 5.0E4, and tolerance set to 0.01 *m/z* or 10 ppm. A smoothing of the chromatograms was applied using Savitsky Golay algorithm, with retention time (RT) smoothing set to 5. The local minimum feature resolver module was used for chromatogram deconvolution, with a chromatographic threshold of 90%, a minimum search range in RT of 0.05 min, a minimum relative height of 1%, minimum absolute height of 5.0E4, minimum ratio of peak top/edge of 2.0, a peak duration of 0.01-1.51 min, and a minimum of 4 data points. The 13C isotope filter module was used by setting a m/z tolerance of 0.01 *m/z* or 5 ppm, RT tolerance of 0.05 min, and setting the maximum charge to 2 and the representative isotope as the most intense one. Then, the features were aligned using tolerance of 0.002 *m/z* or 5 ppm, and RT tolerance of 0.05 min, giving a weight for m/z and RT as 3 and 1, respectively. Gap filling was then performed using an intensity tolerance of 20%, mass tolerance of 0.0015 *m/z* or 5 ppm, RT tolerance of 0.1 min, and a minimum of 1 data point. A features list filter was applied to validate the 13C isotope pattern with a m/z tolerance of 0.0015 *m/z* to 5 ppm, with maximum charge of 2. Then, a duplicate peak filter was applied using a new average filter mode with an *m/z* tolerance of 0.0008 *m/z* or 1.5 ppm and an RT tolerance of 0.1 min. A peak area feature quantification table was exported as a .csv file, in addition to a consensus MS/MS spectral data in .mgf format. Results from MZmine output were used to analyze total ion count for each sample analyzed.

### LC-TIMS-MS Analysis

PEG1000 was analyzed on a Bruker timsTOF PRO 2 mass spectrometer coupled to an Agilent HPLC system to collect ion mobility (IM) data. 2 μL of each standard at 0.1 mg.mL^-1^ in 50:50 MeoH:water (v/v) was injected onto a polar C18 column (Kinetex 2.6 µm Polar C18 100 Å, LC Column 50 x 2.1 mm) at a flow rate of 0.5 mL.min^-1^ held at 40°C. Mobile phase A (99.9% water + 0.1% formic acid) and mobile phase B (99.9% ACN + 0.1% formic acid) were operated with following gradient: t= 0 min, 5 % B; t= 0.5 min, 5 % B; t= 3.0 min, 95 % B; t= 3.5 min, 95 % B; 3.6 min, 5 % B; 5.0 min, 5 % B. ESI was performed in positive ionisation mode with the following parameters: nebulizer gas pressure at 2.2 Bar, dry temperature at 220°C, gas flow 10 L.min^-1^, and spray voltage at 4500 V. MS scan range was set to 20-1200 *m/z* and the following TIMS parameters operated with PASEF (Parallel Accumulation-Serial Fragmentation) were applied: 100 ms ramp time, 100 ms accumulation time, 0.1 - 1.5 Vs.cm^-2^ K0^-1^ start to end range. Five replicates were collected in a randomised order. Raw data is available in MassIVE, ID:MSV000102755.

### TIMS-MS Data Processing

Raw (.d) files were processed with MZmine4 (version 4.10.6). Chromatogram builder parameters were set to a minimum number of consecutive scans of 4, both minimum intensity for consecutive scans and minimum absolute height set to 2.0E2, and a scan-to-scan *mz* tolerance of 0.005 *m/z* or 20 ppm. IMS expander parameters were set as follow: *m/z* tolerance at 0.005 *m/z* or 20 ppm, with the raw data instead of thresholded value set to 10. Smoothing was performed using the Savitzky Golay algorithm set to 5 scans for retention time width, and 5 scans for mobility width. The local minimum feature resolver module was used for chromatogram deconvolution, with a chromatographic threshold of 68.8%, a minimum retention time search range of 0.05 min, a minimum relative height of 0%, minimum absolute height of 2.0E2, minimum ratio of peak top/edge of 1.8, a peak duration of 0.00-1.51 min, and a minimum of 4 data points. The local minimum feature resolver module was also used for mobility deconvolution, with a chromatographic threshold of 90.0%, a minimum mobility search range of 0.03 min, a minimum relative height of 0%, minimum absolute height of 2.0E2, minimum ratio of peak top/edge of 2.5, a peak duration of 0.00-0.30 min, and a minimum of 5 data points. The 13C isotope filter module was used by setting a m/z tolerance of 0.0015 *m/z* or 3 ppm, RT tolerance of 0.04 min, and setting the maximum charge to 15 and the representative isotope as the most intense one. The isotopic peaks finder module was selected with the same parameters as the 13C isotope filter. Features were aligned with an *m/z* tolerance of 0.004 *m/z* or 8 ppm, weighting for *m/z*, RT, and mobility were set to 3, 1, and 1 respectively, RT and mobility tolerances set to 0.1 and 0.01 respectively, and peaks requiring the same charge state. The feature finder module was selected with an intensity tolerance of 20 %, *m/z* tolerance of 0.005 *m/z* or 20 ppm, and RT tolerance of 0.1 mins with a minimum of 2 scans. Duplicate peaks were filtered using the ‘OLD AVERAGE’ mode with an *m/z* tolerance of 0.0008 or 1.5 ppm, a RT tolerance of 0.04 mins, and a mobility tolerance of 0.008. Correlation grouping was carried out with an RT tolerance of 0.06 mins, a minimum feature height of 1.0, and an intensity threshold for correlation of 5.0E2. Feature shape correlation was selected with a minimum number of data points set to 5, and min data points on edge set to 2 with a minimum feature shape correlation of 85 % with PEARSON. Feature height correlation was also selected with minimum samples set to 2, and minimum correlation of 70 % with PEARSON. The feature table (.csv) was exported into Claude for Science for analysis before processing and figure generation in R studio (version 4.5.3).

### NMR Data Analysis and Data Processing

#### Chemicals

All chemicals and reagents used for ^1^H NMR spectroscopy analysis were of analytical grade. Dipotassium phosphate (99%, K_2_HPO_4_), disodium hydrogen phosphate (99%, Na_2_HPO_4_), deuterium oxide (99.9%, D_2_O), and phosphoric acid (≥ 85 wt.% in H_2_O, H_3_PO_4_) for phosphate buffer preparation were purchased from VWR (Radnor, PA, USA). The 3-(trimethylsilyl)propionic-2,2,3,3-*d_4_* acid sodium salt (99%, TSP) was purchased from Sigma-Aldrich (St. Louis, MO, USA). Ultrapure water was generated using a Millipore Direct-Q® 3 UV Water Purification System (Millipore Corp., Bedford, MA, USA)

#### Fecal metabolomics

A total of 867 fecal samples were analysed. For each sample, approximately 200±20 mg of fecal material was weighed into microtubes and stored at −80 °C until analysis. To each sample, 1 mL of ultrapure water was then added, vortexed for 2 min to form a slurry and centrifuged at 4 °C and 24,000 × g for 15 min. Following centrifugation, 630 µL of the supernatant was transferred into a separate microtube and mixed with 70 µL of phosphate buffer in D_2_O (pH 7.4; 1.5 M K_2_HPO_4_/1.5 M NaH_2_PO_4_, 5 mM TSP, and 0.2% NaN_3_). The mixtures were vortexed again for 1 min to ensure complete mixing and centrifuged at 4 °C and 24,000 × g for an additional 10 min. Finally, 600 µL of the resulting supernatant was carefully transferred into 5 mm NMR tubes (Norell Inc., Morganton, NC, USA) and the spectra were immediately acquired within 24h from preparation.

All ^1^H NMR spectra were acquired using a Bruker AVANCE III HD 500 MHz spectrometer operating at 500.11 MHz and equipped with a 5 mm BBFO probe-head (Bruker BioSpin GmbH & Co. KG, Ettlingen, Germany). Spectra were recorded at 298 K using the standard 1D NOESY pulse sequence with water presaturation (Bruker pulse library: *noesypr1d*), employing 65k data points, a spectral width of 16.02 ppm, relaxation delay of 1 s, acquisition time of 4.089 s, a mixing time of 100 ms, receiver gain of 8, 16 dummy scans and 128 scans. Automatic tuning, matching, locking, shimming, and pulse calibration were performed using Bruker standard routines for each sample. All spectra were referenced to the internal standard TSP at 0.0 ppm and manually phased in TopSpin 3.6.4 (Bruker BioSpin).

Sample preparation and spectral acquisition were randomised to minimize potential batch-related effects. Spectra quality was assessed based on the symmetry of the TSP signal and the peak width of less than 1 Hz.

Fecal spectra were pre-processed using an in-house MATLAB^®^ R2020a (The MathWorks, Natick, MA, USA) script, including multipoint baseline correction performed uniformly across all spectra. Spectral regions between δ 0.5–9.0 ppm were segmented into predefined bins based on peak shape, while the residual water region (δ 4.6–5.1 ppm) was excluded from analysis^3^. PEG, δ 3.6–3.8 ppm, was integrated and included the evaluation of the PEG signal.

PEG exposure was assessed by inspection of the δ 3.6–3.8 ppm region, where PEG produces a characteristic broad singlet corresponding to the methylene protons of the ethylene oxide repeating unit (–OCH₂ CH₂ –). Samples were classified as PEG+ by NMR if the bin intensity exceeded a threshold defined manually based on superimposed spectra inspection. PEG+ by NMR and PEG+ by LC-MS were then compared for agreement.

### Metagenomics data analysis

Raw metagenomic reads were pre-processed with Cutadapt and samples from all timepoints for the same patient were co-assembled using MEGAHIT.^4,5^ We produced metagenome-assembled genomes (MAGs) by binning with CONCOCT and MetaBAT2 followed by consensus bin refinement of ambiguous contigs based on coverage profile correlation, and selected the optimal bin representative by maximizing a quality score calculated using a set of 36 SCGs: completeness - 5*contamination.^6,7^ MAGs were pooled and dereplicated at the species level (95% average nucleotide identity) using Galah to obtain species-level genome clusters.^8^ Representative species MAGs were annotated taxonomically using GTDB release 232.^9^

For each dereplicated MAG, we used the same 36 single-copy core gene set and estimated abundance by mapping raw reads from each sample to these ortholog marker genes. Reads that mapped to more than one SCG were discarded and the median value per sample for each MAG across all SCGs was selected as the representative coverage for that MAG after removing outliers identified using median absolute deviation. Differences in community structure from the profile data were assessed using Bray-Curtis dissimilarities visualized by NMDS. PERMANOVA with 9,999 permutations were performed after stratifying for patient. To identify species-level MAGs associated with the presence of PEG, we modeled the centered log ratio (CLR)-transformed relative abundance for each MAG. We used linear models to predict the transformed relative abundances with presence/absence of PEG and patient ID as fixed effects: CLR abundances ∼ PEG + patient. After Benjamini-Hochberg false discovery rate correction for multiple testing, species-level MAGs were ranked by significance and effect size.

### Metabolomics analysis of AIRWAVE cohort

Urine and plasma samples were obtained from the AIRWAVE (AW) cohort - an epidemiological study of microwave radiation exposure from terrestrial trunked radio (TETRA) used by police personnel within the UK.^10^ Urine samples were collected within various clinics local to participating UK police forces, aliquoted into 2 mL cryovials, shipped to a central laboratory at 4 °C within 24 h, and then placed at −80 °C and later at −180 °C in liquid nitrogen for long-term storage. The maximum duration of storage was eight years, and during this time, samples were never thawed. No preservatives were added to the urine samples of either set. Blood samples were collected in the same visit.

Metabolic profiling analysis of urine samples from the AIRWAVE study by both reversed phase liquid chromatography with mass spectrometry detection (RP-LC-MS) (n=1040) and Hydrophilic interaction chromatography with mass spectrometry detection (HILIC-LC-MS) (n=990) revealed intra-sample correlations between mass spectrometry features that were annotated as polymers of PEG and its primary mammalian metabolite PEG-COOH.^11^ These were matched to 990 plasma samples that were collected during the same volunteer visit and were profiled using an adaptation of the HILIC-LC-MS method described in Lewis et al., where the metabolite extraction step was optimised for use with plasma samples, however analytical conditions mirrored the urine analysis exactly.

Raw data sample processing, feature selection and data quality control protocols, were also previously described.^11^ In brief, feature peak picking and alignment was completed using Progenesis QI 2.1 (Nonlinear Dynamics, Newcastle, UK). Feature quality control (QC) checks were completed to remove instrument signal noise. Here, features required a linear response to dilution (r > 0.7) and a relative standard deviation between repeat analyses of an identical QC pool had to be < 30 %.

### Design of PEG MassQL query

PEG ions were searched in public metabolomics data repositories including GNPS/MassIVE and Metabolomics Workbench, which consisted of over ∼3.23 million mass spectrometry files when searched in March 2026. Mass Spec Query Language (MassQL) enables the searching of mass spectrometry data to retrieve MS/MS spectra that contain defined data patterns, and this can also be achieved at the repository level. This search was initially conducted with reference library spectra in the GNPS/MassIVE to test selectivity and to determine the false discovery rate. The GNPS public spectral library contained 2,003,180 MS/MS reference spectra of a wide variety of compounds. A false discovery rate (FDR) was estimated by checking the retrieved spectra for each query that matched. In some cases, reference library compounds had contaminated PEG ions, and these were not accepted as false positives. Our final query had zero false discovery rate. The query was developed to detect PEG oligomers, based on MS2 spectral patterns, consisting of the oxonium ion series at *m/z* 89.0597, 133.0859, and 177.1121, which showed significant intensity with the presence of 221.1383, and 265.1645 ion species. We directed our searches to detect these ions using the following query:

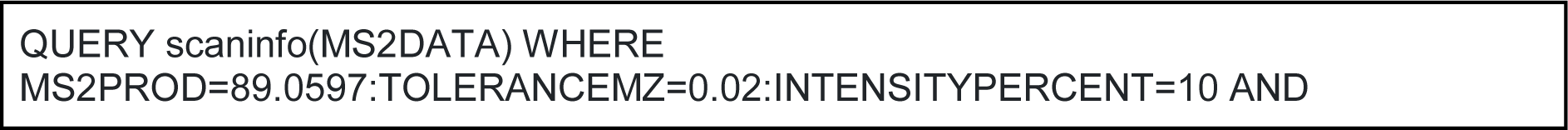
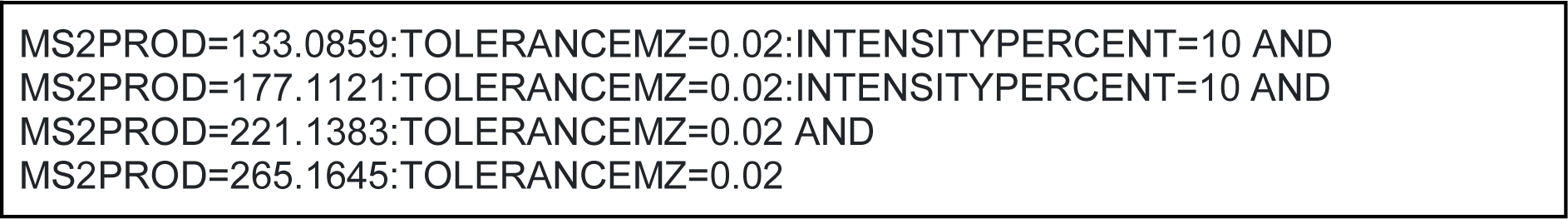

Since 90% of the public metabolomics data was acquired in positive mode, the MassQL searches were mined on data collected in positive ionization mode. The query resulted in a retrieval of >4.4 million spectra from >70,000 mass spectrometry files.

### PEG oligomer search in Personal care products and food items

To test the presence of PEG in personal care products and food items, we conducted a repository scale search of PEG oligomers with cosine similarity above 0.7, minimum matched fragments as 4, and precursor ion and fragment ion tolerances as 0.02 Da. (1) Personal care product MASST: merges the FAST spectral matches across 500 LC-MS/MS files of personal care products (unpublished, https://masst.gnps2.org/personalcaremasst/). (2)foodMASST: merges FASST matches against ∼3,500 LC-MS/MS files of foods and beverages categorized within a food ontology, collected as part of the Global FoodOmics project. These domain-specific MASSTs enabled us to map the distribution of PEG oligomers in personal care products and food items.

### PEG library generation

Once the MassQL jobs with the defined queries were complete, the resulting MGF files were obtained from four jobs: Metabolomics Workbench (https://gnps2.org/status?task=76eebe2be1a54088be028da1bf28a2af), MetaboLights (https://gnps2.org/status?task=798c50ed50984b578d550d5a729b3db0), MassIVE-Orbitrap (https://gnps2.org/status?task=02d9433450fb444caa16a7ef3393b273), and MassIVE-QE (https://gnps2.org/status?task=f1e8b05483b744c09ee81db7c4f53af0). These files were subsequently merged into a single unified MGF file, consolidating all spectral information required for downstream processing.

To ensure data integrity, duplicate scans arising from the overlap between the QE and Orbitrap collections from MassIVE were identified and removed. Spectra were then subjected to precursor mass validation by discarding scans without a precursor mass value, and scans in which the precursor mass did not match the one set for in the MassQL search query. A charge state of +1 was appended to all spectra following this step.

An annotation table in TSV format was generated from the filtered MGF, providing scan-level metadata for subsequent processing steps. Spectra were then integrated with the ReDU sample metadata to enable sample-level filtering. Scans originating from negative ionization modes (electrospray ionization negative, atmospheric pressure chemical ionization negative, and electron ionization), data-independent acquisition (DIA-SWATH) experiments, and a manually curated list of datasets (MTBLS2295, MTBLS2274, MTBLS140) were filtered out.

Next, delta mass values were calculated and retained only when observed in at least two independent datasets. For this step, a core mass of 89.0597 Da was used. Finally, spectra with a precursor ion mass below the highest diagnostic ion of the core structure (minimum precursor mass threshold: 265 Da) were excluded.

The resulting merged MGF file consisted of 4,085,996 scans, with precursor masses ranging from 221.1373 – 1620.9956 Da. All the steps described here were done using the GNPS2 workflow massql_library_generation (task ID https://gnps2.org/status?task=aafeb1057496452a92eb0929deab2807). In order to process this amount of data using a clustering algorithm, the resulting MGF file was separately processed so the scans could be sorted in ascending order of precursor mass value, then split into 5 smaller files.

The resulting spectral files were clustered using Falcon version 1.3 (minimum cluster size: 2 samples; DBSCAN epsilon: 0.1; minimum m/z: 50; minimum peaks per spectrum: 4; minimum m/z range: 1) to group related spectra into representative clusters.^12^ After this first step, the resulting 5 files were clustered once again using the same parameters to reduce redundancy further, resulting in a library with 85,854 representative scans. We also used classical molecular networking to remove all the MS/MS spectra of sub-networks where at least one node contained annotations other than PEG,^13^ reducing the total number of spectra to 85,484. The final library is available at https://library.gnps2.org/api/libraries/GNPS2-PEG-MASSQL-PROPOGATED/mgf.

---

**Figure S1:**
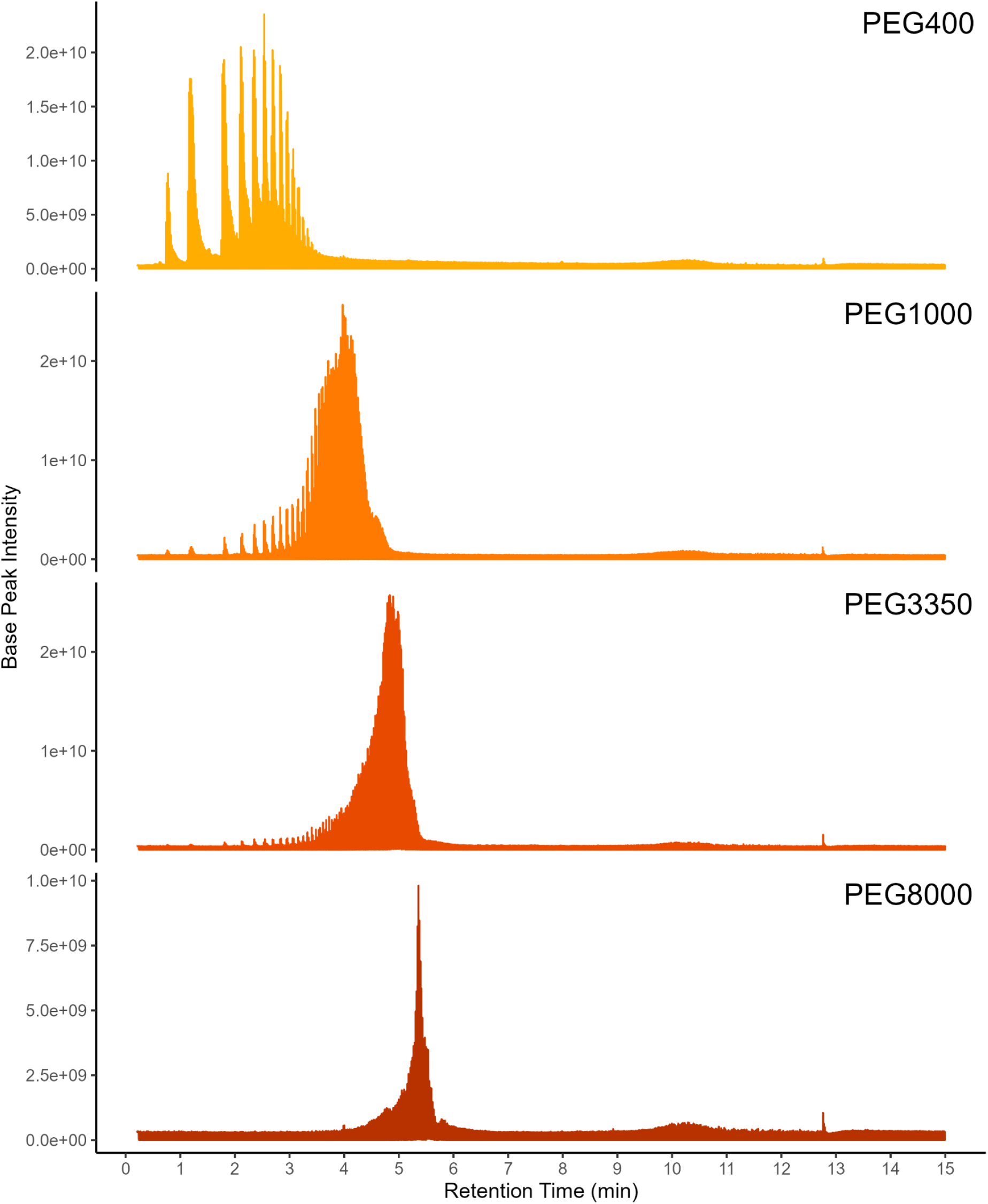
Polyethylene glycol standards elution chromatography in reverse phase LC-MS/MS analysis. Commercially available PEG standards PEG400, PEG1000, PEG3350 and PEG8000 were dissolved in 50:50 MeOH:water at 0.01 mg/mL and analyzed using reverse phase LC-MS/MS analysis using Thermo Orbitrap Q-Exactive.

**Figure S2:**
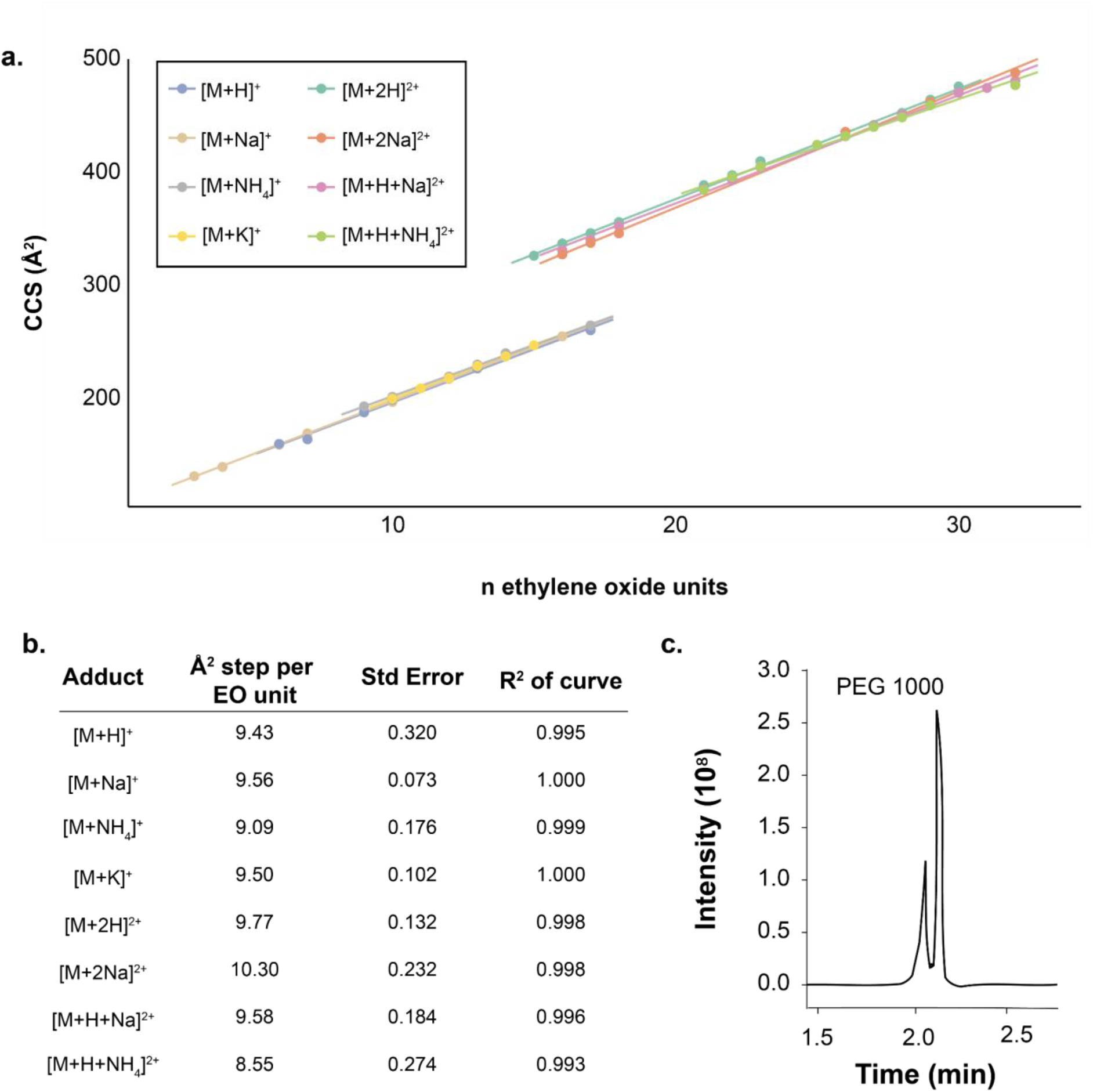
Polyethylene glycol (PEG) collisional cross sectional (CCS) values increase incrementally with additional ethylene oxide (EO) subunits. (a) PEG1000 CCS values increase linearly with additional EO subunits for all detected adduct forms. (b) Linear regression statistics for each detected adduct form of PEG1000. (c) Total ion chromatogram of PEG1000.

**Figure S3:**
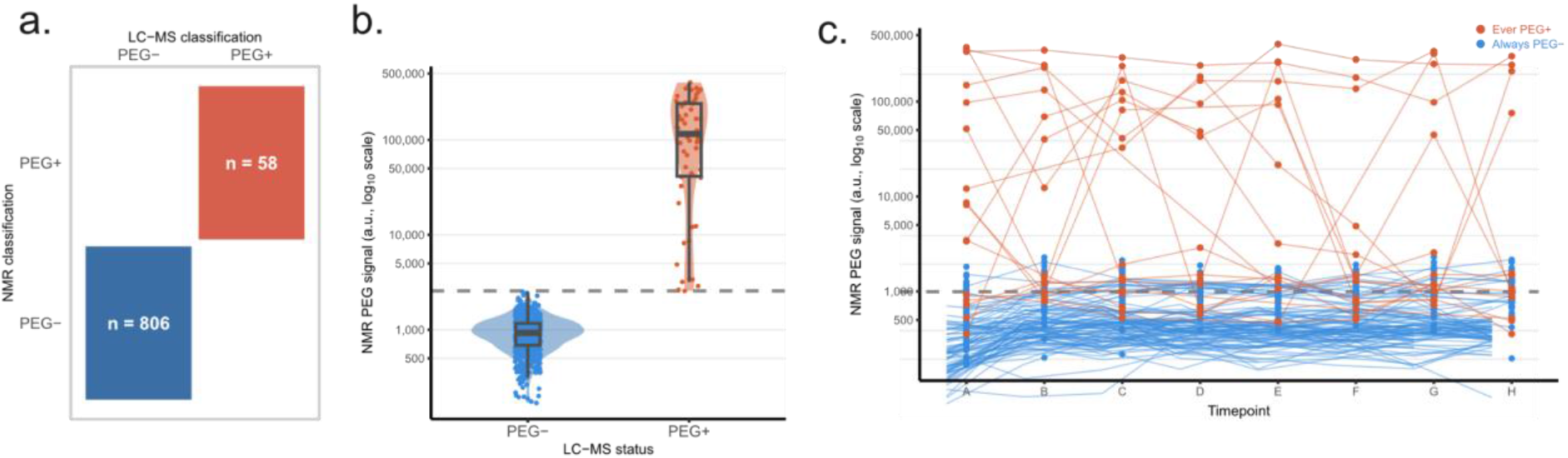
Cross-platform detection of PEG exposure in iPENS fecal samples by NMR spectroscopy and LC-MS/MS. (a) Distribution of NMR PEG signal intensities (integrated intensity of the δ_H_ 3.6–3.8 ppm bin) stratified by LC-MS/MS PEG detection. (b) Contingency table showing agreement between independent NMR- and LC-MS/MS-based PEG detection across all iPENS samples. NMR and LC-MS/MS classifications were performed independently by separate analytical platforms. (c) Longitudinal NMR PEG signal trajectories for individual subjects across study timepoints (A–H),The dashed line indicates the classification threshold (2567.15 a.u.). Orange lines and points indicate subjects with at least one LC-MS/MS PEG+ sample; blue lines indicate subjects consistently classified as PEG-.

**Figure S4:**
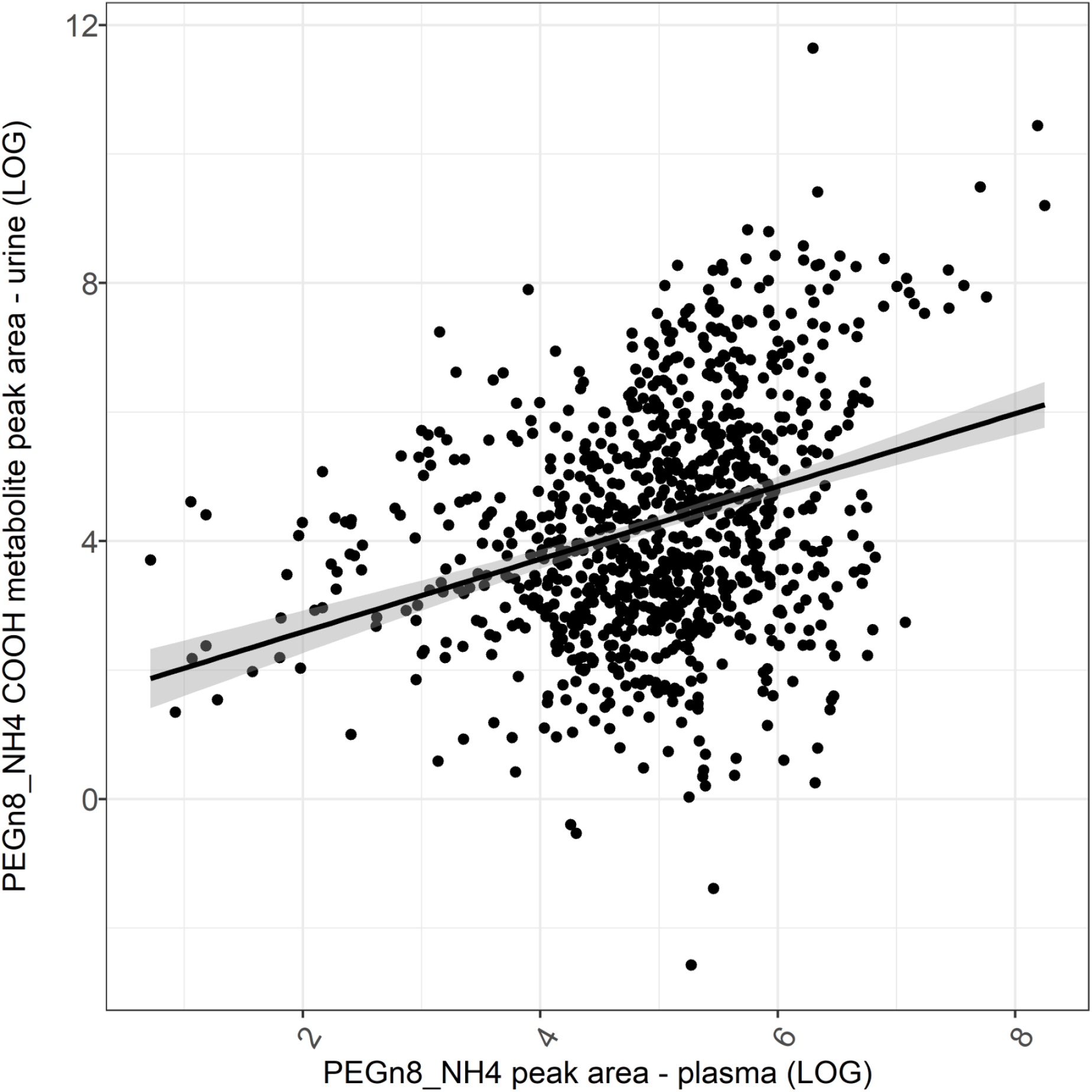
Relationship between circulating plasma PEG and excreted urinal PEG-COOH metabolite. The plot shows data from the AIRWAVE sample cohort (n=990, HILIC-MS urine). The plot shows peak areas (LOG) representative of PEG(n8) in plasma plotted against peak areas (LOG) representative of oxidised PEG-COOH(n8) molecules in matched urine samples from the same study participants. Urine and plasma samples were collected on the same AIRWAVE study visit. Urine metabolite data was corrected for urine volume using NMR creatinine concentration values. Pearson correlation demonstrated that increases of PEG(n8) in plasma have a positive correlation (r = 0.32, *p*<2.2e^-16^) with the PEG-COOH metabolite in urine.

**Figure S5:**
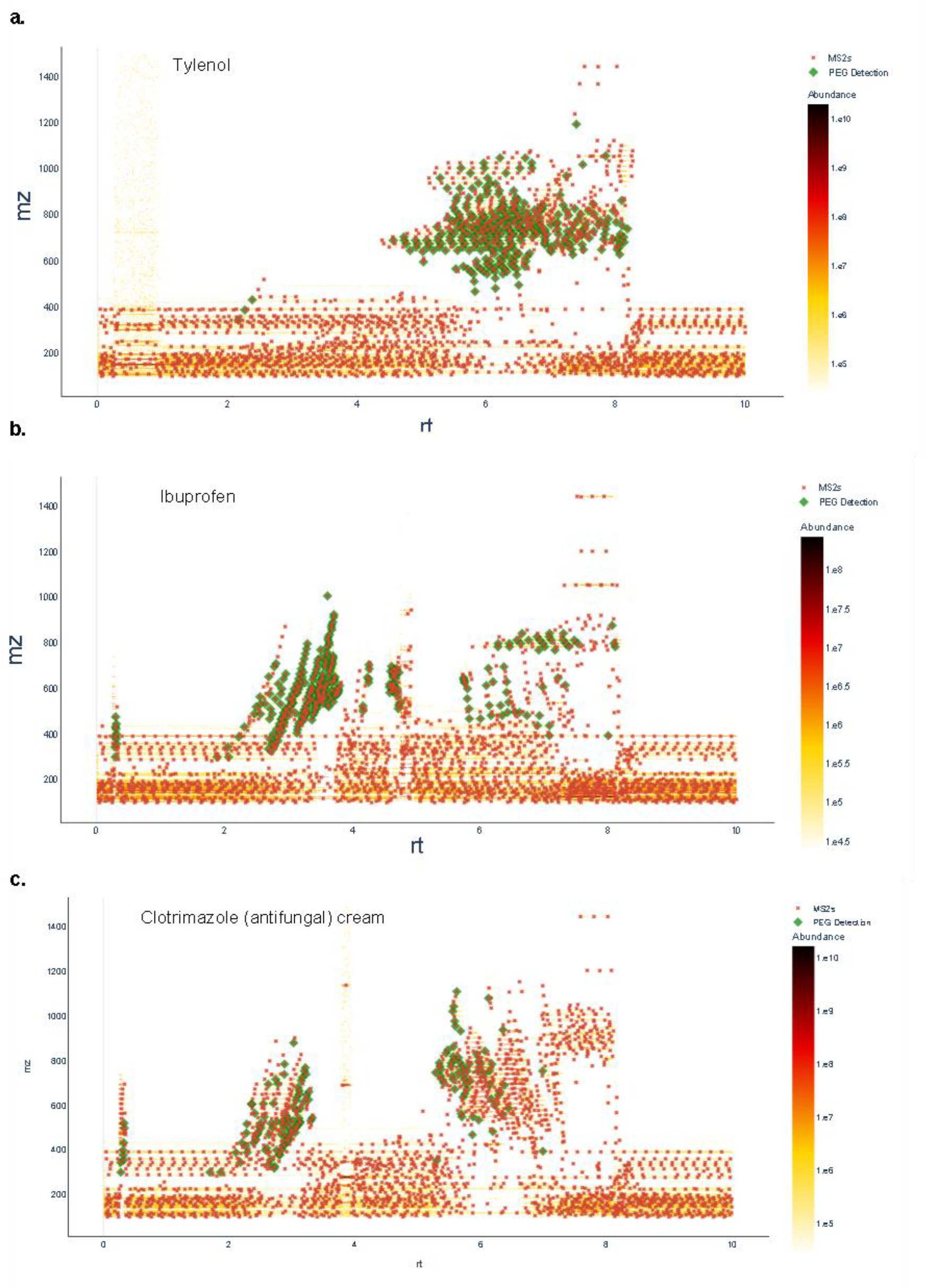
Polyethylene glycol detection in over the counter drug medications. Over the counter drug medications extracted in 100% methanol at 1 mg/ml concentration was analysed using reversed phase LC-MS/MS analysis using thermo orbitrap Q-Exactive. MS2 features highlighted in green, contain diagnostic PEG oxonium ion series *m/z* 89.0597, 133.0859, 177.1121, 221.1383, and 265.1645

**Figure S6:**
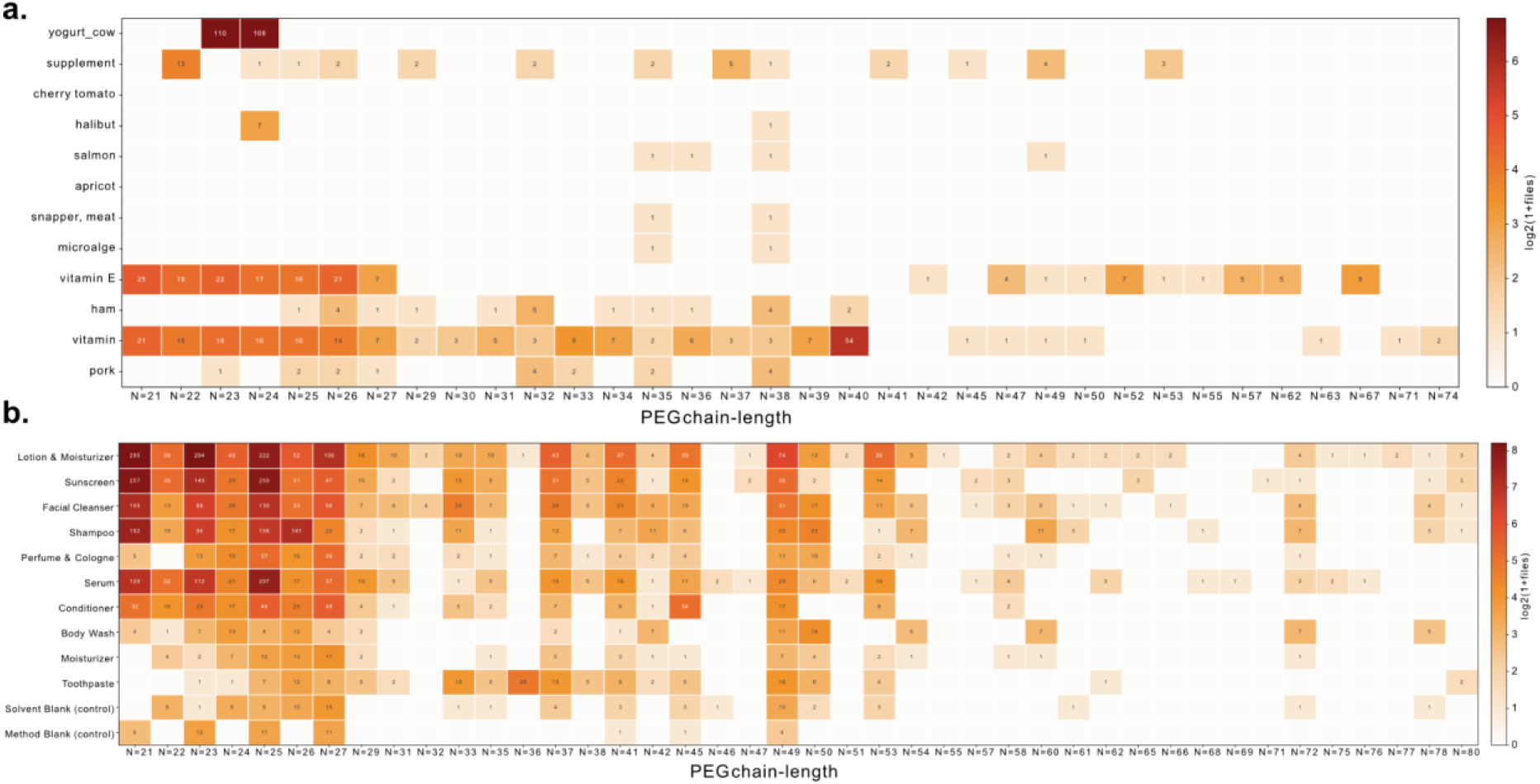
Polyethylene glycol detection in Food and personal care products. High molecular weight (n>20) candidate polyethylene glycol spectra distribution across 3500 food and 500 personal care products analyzed using LC-MS/MS methods.

